# The Effects of Different Fatigue Types on Action Anticipation and Physical Performance in High-level Volleyball Players

**DOI:** 10.1101/2023.11.26.568694

**Authors:** Ying Yu, Liqing Zhang, Ming-Yang Cheng, Zhijun Liang, Ming Zhang, Fengxue Qi

## Abstract

This study examined the effects of different fatigue types on action anticipation and physical performance in high-level volleyball players. Twenty-four participants underwent four counterbalanced conditions: 60-min of cycling at 60% peak power output, 60-min Stroop task, 60-min cycling at 60% peak power output with Stroop task and 60-min neutral documentary to induce physical fatigue (PF), mental fatigue (MF), dual fatigue (DF) and control group (CG), respectively. Action anticipation (anticipation test and visual search test) and physical performance (countermovement jump, T-test, and spike test) were conducted at baseline, immediately after (Post1), and 40-min after fatigue intervention (Post2). DF and PF led to significantly lower jump height, T-test completion time, and spiking speed at Post1 and Post2 compared with CG and MF. Compared with PF, DF led to a significant decline in jumping, agility, and spiking performance at Post1 and decreased jumping performance at Post2. MF significantly decreased reaction time in the anticipation test at Post1 compared with CG. Reaction times in the visual search test were significantly lower with DF and MF at Post2 compared with CG. PF led to a greater decline in physical performance and this was exacerbated in DF, yet anticipation performance remained lesser impact.

## Introduction

Volleyball is a complex team sport with an average match duration of 60–90 min, imposing distinct physical and cognitive demands (Formenti et al., 2020). As an open-skill sport, volleyball requires motor actions characterized by efficient visual search and action anticipation in dynamic and intricate settings (Nuri et al., 2013). A heightened ability for action anticipation is closely associated with the skill of effectively “reading the game” and possessing superior game intelligence (Williams et al., 2011), which is a prerequisite for executing techniques and tactics with success (Nuri et al., 2013). Consequently, physical performance, including elements such as vertical jumping, agility to cover short distances, and spiking (Plesa et al., 2021), as well as perceptual–cognitive performance, are key determinants of volleyball performance. Fatigue is prevalent in sports, particularly in high-intensity and in prolonged activities like volleyball (Knicker, Renshaw, & Cairns, 2011; Tsarbou et al., 2021). A comprehensive understanding of how fatigue affects action anticipation and physical performance is crucial for devising effective training strategies for high-level volleyball players.

Fatigue is a multifaceted symptom that affects various aspects of the body, encompassing cardiorespiratory, neuromuscular, physiological, and cognitive functions (Bestwick-Stevenson et al., 2022; Knicker, Renshaw, Oldham, et al., 2011; Pageaux & Lepers, 2018). It can be classified into different types, each with unique characteristics. Physical fatigue (PF) is a physiological phenomenon marked by decreased neural excitability in the motor cortex, leading to an inability to sustain the necessary force output following prolonged physical activities (Thomas et al., 2017). PF diminishes neuromuscular capacity, primarily affecting strength, speed, and endurance (Lin et al., 2021; Marcora et al., 2008; Skurvydas et al., 2011). Additionally, PF has been associated with impaired anticipation and decision-making abilities, possibly associated with reduced activation in the prefrontal cortex (D. B. Alder et al., 2019; Parkin et al., 2017). Mental fatigue (MF) is a psychological state induced by prolonged cognitive activity or high cognitive demand, resulting in a feeling of exhaustion or weariness (Marcora et al., 2009). MF affects the activation of the prefrontal cortex, reducing attentional resources and, consequently, affecting cognitive function, technical abilities, and tactical execution (Filipas et al., 2021; Kosack et al., 2020; Smith et al., 2016). Numerous studies have demonstrated the detrimental effects of MF on endurance (Holgado et al., 2021; Penna et al., 2018), training volume (Queiros et al., 2021) and balance (Noe et al., 2021). However, its influence on neuromuscular performance, such as explosive power and maximal strength, appears to be negligible (Duncan et al., 2015; Pageaux et al., 2015).

Many sport-related and non-sport-related situations impose high cognitive demands, which can lead to MF (Thompson et al., 2021). Extended training sessions or competitions are more likely to lead to the accumulation of physical and mental loads. There are common neuromodulatory pathways between PF and MF (Mehta & Parasuraman, 2014). The combination of PF and MF, known as dual fatigue (DF), can exacerbate the inhibition of prefrontal cortex activity and result in more severe impairments compared with the negative effects of single type of fatigue (D. Alder et al., 2021; Dailey et al., 2015; Kelly et al., 2012). Barzegarpoor et al. reported that, in male recreational cyclists, DF negatively affected endurance more than PF did, with no significant differences observed in heart rate or maximum voluntary contraction (Barzegarpoor et al., 2020). The impairment caused by DF appears to be primarily psychologically rather than physiologically mediated. However, Rubio-Morales et al. found that, among physically active university students, only MF resulted in significant impairments in reaction time following exposure to DF, MF, and PF (Rubio-Morales et al., 2022). A plausible explanation for these inconsistent results is that fatigue may manifest differently among athletes participating in various sports disciplines and at different levels.

Open-skill sports demand that athletes respond to unpredictable environments, demanding enhanced visual attention, precision in execution, and decision-making abilities (Bishop et al., 2013; Kim et al., 2008; Nakamoto & Mori, 2008). However, the effects of different fatigue types on high-level athletes may have different characteristics (Coyne et al., 2021). Parkin et al. reported that PF can enhance rather than impair decision-making performance in top athletes (6 Olympic medalists) (Parkin et al., 2017). High-level athletes demonstrate higher resistance to fatigue (Kosack et al., 2020), and they consistently exhibit superior physical and cognitive performance compared to novice and younger individuals (De Waelle et al., 2021; Piras et al., 2014; Trecroci et al., 2021). Understanding the effects of different fatigue types on action anticipation and physical performance can help in designing targeted training interventions for high-level volleyball players. However, no existing research has delved into the distinct effects of different types of fatigue on action anticipation and physical performance in high-level volleyball players.

The primary objective of this study was to investigate the effects of different types of fatigue on action anticipation and physical performance among high-level volleyball players. As noted by Thomas et al., the effects of prolonged exercise on the central nervous system were primarily associated with exercise duration, with intermittent and continuous exercise yielding similar outcomes (Thomas et al., 2017; Thomas et al., 2016; Thomas et al., 2015). To induce distinct types of fatigue, we employed three methods: 60 min of cycling at 60% peak power output (PPO), a 60-min Stroop task, and 60 min of cycling at 60% PPO combined with the Stroop task to induce PF, MF, and DF, respectively. We hypothesized that both action anticipation and physical performance would be reduced in DF.

## Method

### Participants

Twenty-four highly trained volleyball players (mean `±` SD age: 21.7 ± 1.2 years, body mass: 77.8 ± 7.78 kg, body mass index: 22.37 ± 2.19 kg/m^2^, experience: 6.37 ± 1.58 years, peak power: 235 `±` 44 W) participated in this study. All participants were recruited from Beijing Sport University, and no dropouts occurred. They brought substantial competitive experience to the research, having competed in national-level events for an average of 6.37 years. The sample size was determined based on data from previous studies (D. Alder et al., 2021) and calculated using G*Power software (version 3.1) with an effect size of f = 0.3, an alpha level of .05, and a power of .80. Allowing for 20% attrition, we recruited 20% more participants than the power calculation indicated were needed. All experimental procedures strictly adhered to the ethical principles set forth in the Declaration of Helsinki and received approval from the ethics committee at Beijing Sport University (2022166H).

### Psychophysiological parameters

The Rating of Perceived Exertion (RPE) was employed to assess participants’ perceived effort (Ribeiro et al., 2008). This scale requires participants to rate the overall effort required to complete a task, ranging between 6 (*no exertion*) to 20 (*maximal exertion*). The 100-mm Visual Analogue Scale (VAS) was used to assess the subjective perception of MF (Dernis-Labous et al., 2003; Kosack et al., 2020). Participants marked how mentally fatigued they were on a scale anchored by *not all exhausted* and *completely exhausted* by placing a mark on a 100-mm line. The VAS is a widely recognized tool for assessing the extent of MF and is known for its high validity in evaluating neural excitability changes during fatigue (Marzouk et al., 2023). The heart rate (HR) was monitored using a sensor (H10; PolarTM, Finland).

### Physical performance

Agility was assessed using the modified T-test (SASSI et al., 2009), involving a combination of forward sprint, lateral slides, and a backward run. Participants initiated the test with a forward sprint covering a distance of 5 m, where they touched a central cone with their right hand. While maintaining a forward orientation and avoiding crossing their feet, they moved 2.5 m to the left, touching a cone with their left hand. Following this, they swiftly slid 5 m to the right, ensuring contact with a cone with their right hand. Subsequently, they reversed direction, gliding back to the left to touch the central cone. The test concluded with participants sprinting backward to the starting line as rapidly as possible. Each participant performed two attempts, with a brief 1–2 min rest between them. The time was measured using timing gates (Wireless Timing Systems, Shiniangong, China). The reliability of the modified T-test was assessed using the intra-class correlation coefficient (ICC; ICC = 0.899).

Jumping performance was assessed using the countermovement jump (CMJ). During this assessment, participants started from an upright standing position with hands on their hips and jumped as high as possible. This involved descending to their usual squat depth, followed by an explosive jump to attain maximum height. The landing was executed in an athletic position. Each participant completed three maximal attempts, with a 30-s interval between each repetition. The CMJ measurements were captured using an optical jump system (Microgate, Bolzano, Italy). The CMJ test showed an ICC of 0.982.

The spike test (ST) was performed according to a previously described protocol (Valades, 2009). In brief, players were tasked with hitting a ball with maximal force toward a target zone (1.5 × 1.5 m). Participants were required to self-toss the ball and execute the spike without jumping or lifting their feet off the ground. A researcher stood to the side to monitor the performance closely. If any aspect of the execution was incorrect, the test was repeated. A Bushnell Radar Gun (Bushnell Outdoor Products, Kansas, USA) was positioned to measure the velocity of the ball within the target zone. Rigorous monitoring of ball pressure, radar orientation, and calibration was undertaken for each test. Each player was granted three trials, and the average speed was recorded. The ST showed an ICC of 0.967.

### Action anticipation

In the anticipation test (AT) (De Waelle et al., 2021), videos were captured at the center of the court, positioned at a height of 1.30 m above the ground, using a high-definition digital camera (Sony A7M4; Sony, Tokyo, Japan) with a frame rate of 50 frames/s. The video clips illustrated a situation in which a free ball was expertly distributed to the setter’s position. Afterward, the setter transferred the ball to one of three conditions: (1) a long backward pass to the outside spiker, (2) a brief forward pass to the central spiker, or (3) an extended forward pass to the other outer spiker (right side). As the play neared the set-up phase, the video was occluded (i.e., a black screen was presented), referred to as “viewing conditions”. Specifically, the videos were intentionally occluded at the moment the ball was received by the setter. Participants were instructed to respond promptly and with precision, identifying which of the three passing options the setter would choose by pressing the corresponding key on the keyboard. To ensure that participants understood the test, they received a comprehensive explanation, which included a familiarization clip and practice trials. During the test, a total of 100 video clips were presented, each lasting about 2–4 s. These clips were displayed in a randomized order for each participant to mitigate any potential order effects. Participants were granted a 5-s window within which to respond. Response times (ATRT) and accuracy (ATACC) were recorded as critical performance measures. The ICC showed reliable results, with mean values of 0.861 for ACCRT and 0.893 for ATACC.

In the visual search test (SVT) (Parkin & Walsh, 2017), participants were required to conduct a visual search within an array to determine the presence or absence of a specific target image—in this case, a red square. The visual array included three different types of distractors: green squares, green circles, and red circles. The groups were divided into 3, 6, and 9 items according to the number of displayed items. Two different trial types were included: 1) the target was present, and the display item featured a red square; or 2) the target was not present, and the display item did not contain a red square. Throughout the test, a total of 156 clips were shown across two blocks. Participants could rest between blocks. Response time (SVTRT) was recorded. The ICC provided strong evidence of reliability, with mean values of 0.912.

### Fatigue protocol

An incremental load test was used to assess PPO (Strasser et al., 2016). Participants began with a 3-min warm-up at 75 W, after which the workload increased by 25 W/min while maintaining a cadence of 60 rpm. Verbal encouragement was continuously provided to the participants until they reached the point of exhaustion. If a participant was exhausted, the cycling power was reduced for active recovery. Heart rate and RPE were collected every minute during the exercise, and PPO was collected immediately after exhaustion. Exhaustion was considered reached by achieving two of the following three criteria: (i) maintaining speed below 60 rpm for 15 s; (ii) reporting an RPE of 17–20; (iii) reaching a heart rate within the range of 90%–100% of the maximum heart rate (207 - 0.7*age).

PF was induced through 60 min of cycling at 60% PPO on a cycle ergometer (Ergoselect 100k, Ergoline, Bitz, Germany) while the subjects watched a neutral documentary titled ‘*Human and Nature.*’ Fatigue was considered reached when the following criteria were met (Barzegarpoor et al., 2020; Ribeiro et al., 2008; Thomas et al., 2016): (i) reporting an RPE above 15, and (ii) HR reached `±` 15% of maximal heart rate of incremental load test.

MF induction consisted of a 60-min computerized version of the Stroop test while sitting on cycle ergometer. In this cognitive task, four words (red, blue, green, and yellow) were displayed one at a time on a 24-inch screen with a black background. Participants were instructed to respond to the selected stimulus using a computer keyboard by pressing one of four keys that matched the color of the word rather than the word’s meaning. For instance, when the word “green” appeared in yellow color, the correct response was to press the key corresponding to the color yellow. The task contained 50% incongruent trials. Each word remained on the screen for 1000 ms, followed by a 1500-ms interval with a black screen before the next word appeared. MF was considered reached when a VAS score above 50 was reported (Dernis-Labous et al., 2003; Kosack et al., 2020).

DF was induced using 60 min of cycling at 60% PPO while performing the Stroop task. DF was considered to be induced when the following criteria were met: (i) reporting an RPE above 15, (ii) reporting a VAS score above 50, and (iii) HR reached `±` 15% of maximal heart rate of incremental load test. The control group (CG) watched a 60-min neutral documentary titled ‘*Human and Nature*’ while sitting on cycle ergometer.

### Procedure

Each participant made a total of five visits to the laboratory, with each visit being spaced at least 7 d apart. Before the experiment, the participants were informed of the experimental procedures and precautions. Informed consent was obtained from each participant, and they were familiarized with the test procedures, signing any necessary consent forms. Twenty-four hours before each visit, participants were instructed to refrain from consuming alcohol or caffeinated beverages (e.g., coffee, tea, cola, energy drinks, caffeinated supplements, and chocolate), ensuring a minimum of 7 h of sleep, avoiding strenuous mental and physical activities, and abstaining from eating at least 2 h prior to testing. Participants were also required to disclose any medication or drug use, as well as to report any acute illness, injury, or infection. The time of testing, environmental conditions, and equipment settings were also standardized.

At Visit 1, before the baseline test, participants engaged in familiarization trials, which significantly improve the reliability of testing, particularly in highly trained individuals (Driller et al., 2013). Wearing earplugs, participants first completed the action anticipation and visual search tests on a computer equipped with a 24-inch screen. Participants then performed a 15-min standardized warm-up, consisting of 8 min of dynamic stretching exercises for the upper and lower limbs to increase core and muscle temperatures and 7 min of specific volleyball exercises. Subsequently, they proceeded to the physical performance tests, including the incremental load power test used to record their PPO.

At Visits 2–5, following a 15-min warm-up, the experiment included four treatment conditions: PF, MF, DF, and CG. The order of the four treatment conditions was counterbalanced across participants to control for order effects. HR, RPE, and VAS data were collected before and after completing the fatigue intervention. Then, participants wore earplugs and completed the visual search and anticipation tests in a counterbalanced order, and then completed the physical performance test using the randomized order (Post 1). The entire testing process took approximately 30 min. Subsequently, after a 10-min rest period, a second post-test (Post 2) was conducted, employing the same methods.

### Statistical Analysis

Data for CMJ, T-test, ST, SVT, and AT were standardized through a baseline transformation (baseline/baseline, Post 1/baseline, Post 2/baseline). Data normality was assessed using the Shapiro–Wilk test. One-way repeated-measures analysis of variance (RM-ANOVA) was applied to RPE, HR, and VAS data to ensure that there were no differences before fatigue interventions between sessions. Within-subjects RM-ANOVA incorporating the factors of session (DF, PF, MF, and CG) and time (baseline, Post 1, and Post 2) was used to compare the effects of different fatigue types on three measures of physical performance (CMJ height, Agility T-test completion time, and spike speed) and three measures of action anticipation (ATRT, ATACC, and SVTRT). Greenhouse–Geisser procedures were used to correct for violations of the sphericity assumption. Subsequently, the post-hoc Fisher’s least significant difference test was performed to identify specific differences on measures of physical performance and action anticipation between conditions. Effect size estimates were calculated in the form of partial eta-squared (η*_p_*^2^) for the ANOVAs and were classified as small (0.010 < η*_p_*^2^ ≤ 0.059), moderate (0.059 < η*_p_*^2^ ≤ 0.138), or large (η*_p_*^2^ > 0.138). Additionally, Cohen’s *d* (effect size) was reported for *Post-hoc* comparisons and was interpreted as small (0.2), moderate (0.5), or large (0.8). All statistical analyses were conducted with a significance level of P < .05 (two-tailed). Statistical analyses and figure creation were performed using SPSS 26.0 (IBM Corp., Armonk, NY, USA) and GraphPad Prism 8.0 (Graph Pad, La Jolla, CA, USA).

## Results

### Fatigue-inducing conditions

Descriptive data on RPE, HR, and VAS are presented in Table 1. There were no significant differences in VAS (P = 0.315), HR (P = 0.171), and RPE (P = 0.872) among the DF, MF, PF, and CG conditions before fatigue interventions. On the other hand, the results suggested that the fatigue tasks successfully induced different types of fatigue.

**Table 1.** RPE, HR and VAS before and after fatigue intervention (Mean ± SD).

| Fatigue type |  | RPE | HR | VAS |
| --- | --- | --- | --- | --- |
| Dual fatigue | Before | 7.20 ± 1.47 | 77.04 ± 10.60 | 15.67 ± 10.30 |
| | After | 19.33 ± 2.03 | 170.37 ± 12.27 | 64.37 ± 20.67 |
| Mental fatigue | Before | 7.29 ± 1.27 | 79.96 ± 6.40 | 15.88 ± 11.71 |
| | After | 9.71 ± 2.84 | 72.62 ± 7.44 | 52.04 ± 19.69 |
| Physical fatigue | Before | 7.37 ± 1.28 | 76.41 ± 6.00 | 17.12 ± 8.50 |
| | After | 19.04 ± 1.23 | 169.41 ± 13.33 | 45.58 ± 22.11 |
| Control group | Before | 7.16 ± 1.43 | 78.25 ± 7.50 | 13.16 ± 10.03 |
| | After | 8.54 ± 2.51 | 76.83 ± 9.59 | 15.29 ± 11.87 |

### Physical performance Countermovement jump

For the CMJ, the RM-ANOVA analysis showed a significant main effect of time (F_2, 46_ = 127.518, P < 0.001, η*_p_*^2^ = 0.847), and session (F_1.649, 37.918_ = 33.485, P < 0.001, η*_p_*^2^ = 0.593), as well as an interaction effect of time and session (F_2.605, 59.905_ = 27.883, P < 0.001, η*_p_*^2^ = 0.548). Post-hoc tests for CMJ indicated a significant impairment at Post 1 in DF (P < 0.001, ES = 0.75) and PF (P < 0.001, ES = 0.69) as well as at Post 2 in DF (P < 0.001, ES = 0.63) and PF (P < 0.001, ES = 0.58) compared with CG. Furthermore, there were significant CMJ impairments at Post 1 in DF (P < 0.001, ES = 0.72) and PF (P < 0.001, ES = 0.63), and at Post 2 in DF (P < 0.001, ES = 0.63) and PF (P < 0.001, ES = 0.58) compared with MF.

Notably, the CMJ performance in DF deteriorated even further when compared with PF at Post 1 (P = 0.004, ES = 0.39) and at Post 2 (P = 0.042, ES = 0.30) (Figure 2A). Significant differences in CMJ height was observed for DF, PF, and MF at Post 1 and Post 2 compared with baseline and between Post 1 and Post 2 (all values of P < 0.006). CG displayed significant CMJ impairments at Post 1 and Post 2 compared with baseline (all values of P < 0.003).

**Figure 1.**
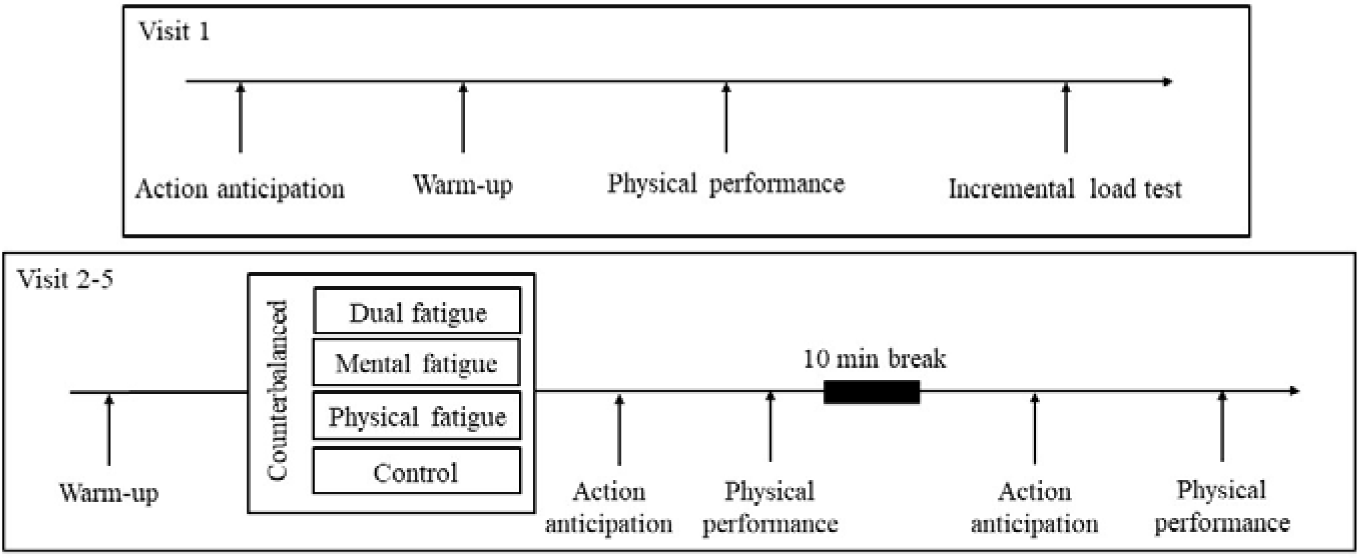
Study protocol of dual fatigue, mental fatigue, control, and physical fatigue. Action anticipation included anticipation test and visual search terst, physical performance included agility T-test, countermovement jump and spike test. Visit 1 included baseline test and incremental load test to record peak power output. Visit 2-5 included four counterbalanced experimental conditions.

**Figure 2.**
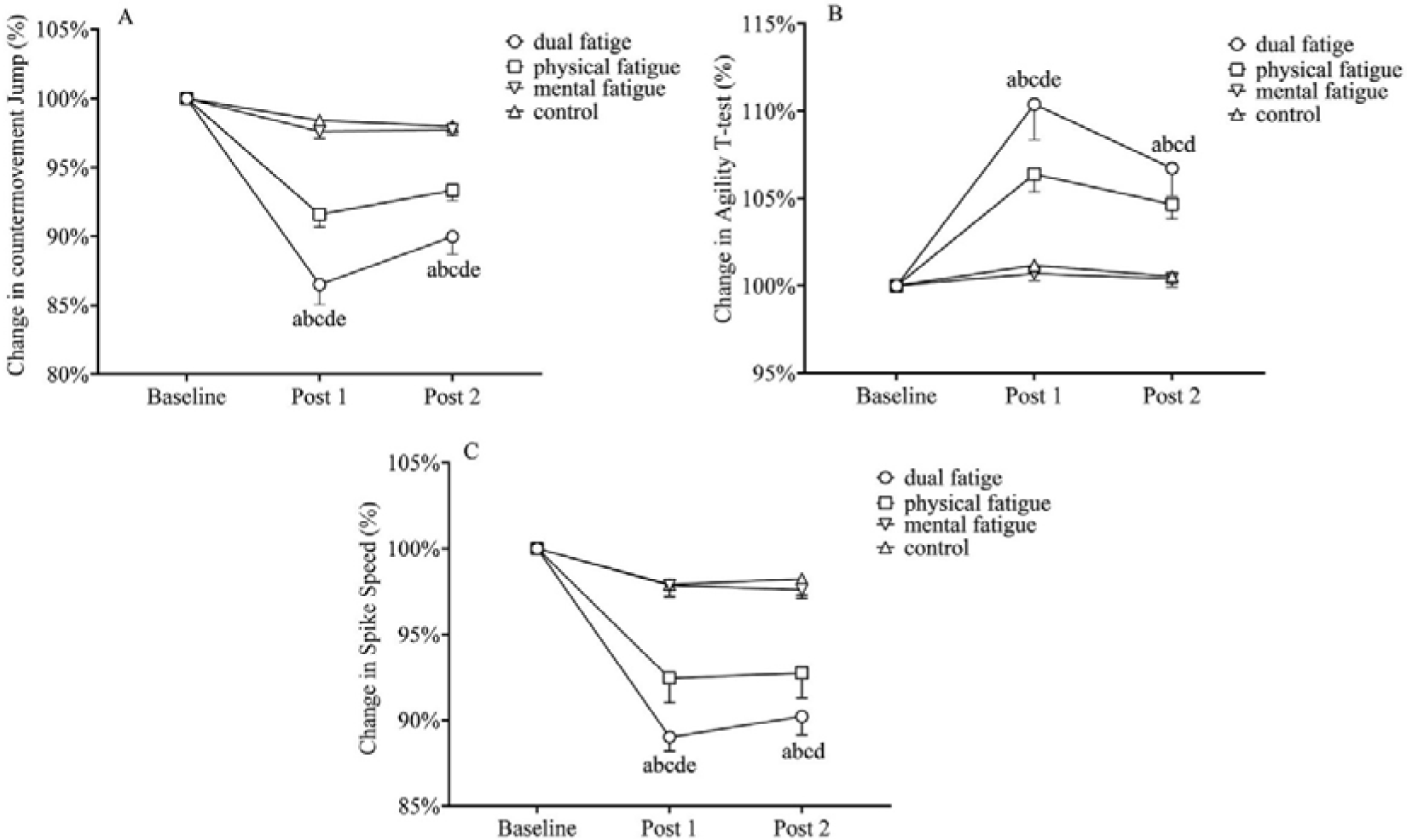
The group data jump height in countermovement jump (panel A), time in Agility T-test (panel B) and spike speed in Spike test (panel C) in immediately after the intervention (Post 1), and 40-min immediately following the intervention (Post 2). a: significant difference from dual fatigue to control. b: significant difference from dual fatigue to mental fatigue; c: significant difference from physical fatigue to control; d: significant difference from physical fatigue to mental fatigue; e: significant difference from dual fatigue to physical fatigue. Level of significance was set at p < 0.05.Error bars represent standard error (SEM).

### T-test

For the T-test, the RM-ANOVA demonstrated a significant main effect of time (F_1.423, 32.740_ = 26.223, P < 0.001, η*_p_*^2^ = 0.553), and session (F1.423, 32.724 = 21.287, P < 0.001, η*_p_*^2^ = 0.481), as well as an interaction effect between time and session (F_2.337, 53.744_ = 16.136, P < 0.001, η*_p_*^2^ = 0.412). Further post-hoc testing revealed that there was a notable decrease in the DF condition for the T-test completion time at Post 1 (P < 0.001, ES = 0.56) and Post 2 (P < 0.001, ES = 0.48) compared with the CG condition. Similarly, PF displayed a significant impairment in T-test completion time at Post 1 (P < 0.001, ES = 0 59) and Post 2 (P < 0.001, ES = 0.54) compared with CG. Additionally, a significant impairment in T-test completion time was observed for DF (P < 0.001, ES = 0.54) and PF (P < 0.001, ES= 0.56) at Post 1 and for DF (P < 0.001, ES = 0.47) and PF (P < 0.001, ES = 0.54) at Post 2 compared with MF. Importantly, in the DF condition, T-test completion time was even further reduced when compared with PF at Post 1 (P = 0.021, ES = 0.24) (Figure 2B). Significant differences in T-test completion time were observed for DF, PF, and MF at Post 1 compared with baseline and between Post 1 and Post 2 (all values of P < 0.045). DF and PF led to significant impairments in completion time at Post 2 compared with baseline (all values of P < 0.001).

### Spike test

For the ST, RM-ANOVA revealed a significant main effect of time (F_1.476, 33.945_ = 73.155, P< 0.001, η*_p_*^2^ = 0.761), and session (F = 24.364, P < 0.001, η*_p_*^2^ = 0.514), as well as an interaction effect between time and session (F_6, 138_ = 14.259, P < 0.0001, η*_p_*^2^ = 0.383). A more detailed examination using post-hoc testing revealed that, for the ST, ball speed was significantly reduced at Post 1 for both DF (P < 0.001, ES = 0.76) and PF (P < 0.001, ES = 0.69) and at Post 2 for DF(P < 0.001, ES = 0.64) and PF (P = 0.002, ES = 0.42), compared with CG. Additionally, ball speed was significantly reduced at Post 1 for both DF (P < 0.001, ES = 0.77) and PF (P = 0.002, ES = 0.44), and at post 2 in DF (P < 0.001, ES = 0.67) and PF (P = 0.004, ES = 0.41) compared with MF. Importantly, in DF, the ball speed was even further reduced when compared with PF at Post 1 (P = 0.011, ES = 0.29) (Figure 2C). Significant reductions in ball speed were observed for DF, PF, and MF at Post 1 and Post 2 compared with baseline (all values of P < 0.03), and significant reductions in ball speed were observed for CG at Post 1 (P = 0.008) compared with baseline.

### Action anticipation

#### Reaction time in anticipation test

Regarding ATRT, RM-ANOVA showed significant main effects of time (F_1.135, 26.109_ = 12.982, P < 0.001, η*_p_*^2^ = 0.361), and session (F3, 69 = 3.243, P = 0.027, η*_p_*^2^ = 0.124), as well as a significant interaction effect between time and session (F_6, 138_ = 2.640, P = 0.019, η*_p_*^2^ = 0.103). More detailed examination using post-hoc testing revealed that ATRT was significantly higher at Post 1 (P = 0.001, ES = 0.30) for MF compared with CG (Figure 3A). Significant improvements in ATRT were observed for DF, PF, and CG at Post 1 and Post 2 compared with baseline (all values of P < 0.019). A significant improvement in ATRT was observed for MF at Post 2 compared with baseline and between Post 1 and Post 2 (all values of P < 0.004).

**Figure 3.**
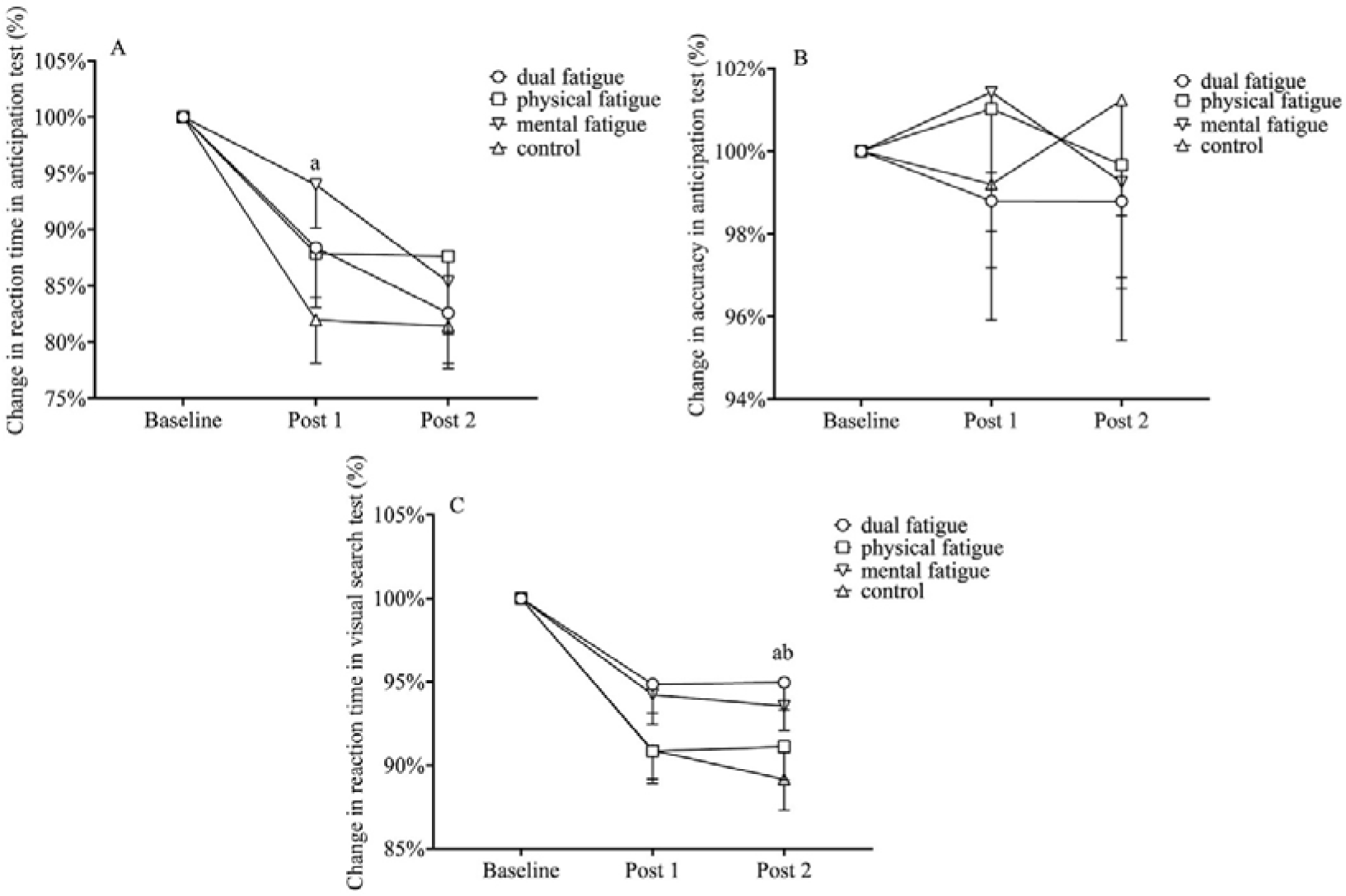
The group data of response time (panel A) and accuracy (panel B) in anticipation and response time in visual search test (panel C) on four sessions in immediately after the intervention (Post 1), and 40-min immediately following the intervention (Post 2). a: significant difference from dual fatigue to control. b: significant difference from mental fatigue to control. Level of significance was set at p < 0.05. Error bars represent standard error (SEM).

### Accuracy in anticipation test

Regarding ATACC, RM-ANOVA revealed no significant main effects of time (F_1.094, 25.163_ = 0.019, P = 0.909, η*_p_*^2^ = 0.020), or session (F2.322, 53.414 = 0.405, P = 0.699, η*_p_*^2^ = 0.017) and no significant interaction effect between time and session (F_6, 138_ = 1.018, P = 0.416, η*_p_*^2^ = 0.042) (Figure 3B).

### Reaction time in visual search test

Regarding SVTRT, RM-ANOVA showed a significant main effect of time (F_1.586, 36.487_ = 27.357, P < 0.001, η*_p_*^2^ = 0.543), and session (F_3, 69_ = 3.816, P = 0.014, η*_p_*^2^ = 0.142), as well as of the interaction between time and session (F_6, 138_ = 2.245, P = 0.042, η*_p_*^2^ = 0.089). Post-hoc tests revealed that SVTRT was significantly higher at Post 2 for DF (P < 0.001, ES = 0.32) and MF (P = 0.019, ES = 0.26) compared with CG (Figure 3C). Significant SVTRT improvements were observed for DF, PF, MF, and CG at Post 1 and Post 2 compared with baseline (all values of P < 0.006).

## Discussion

This study aimed to investigate the effects of different fatigue types on action anticipation and physical performance in high-level volleyball players. The main findings of this study showed that DF and PF significantly reduced jump height and led to a decrease in the T-test completion time and spike speed compared with MF and CG. DF led to significantly decrease jump height, T-test completion time, and spike speed at Post 1 and CMJ height at Post 2 compared with PF. Conversely, MF did not lead to any significant effects on these parameters. Notably, neither DF nor PF had a significant effect on action anticipation in high-level volleyball players compared with the CG condition. MF led to significantly decreased reaction times in the anticipation test at Post 1 compared with CG. DF and MF also led to significantly decreased reaction times in the visual search test at Post 2 compared with CG.

PF had a negative impact on jumping, agility, and spiking performance when compared with MF and CG. These results were in accordance with a study by Lin et al., which highlighted the adverse effects of PF on jumping ability and landing strategy in volleyball players (Lin et al., 2021). Skurvydas et al. showed that PF decreased maximum voluntary contraction and the central activation ratio in long-distance runners, sprint runners, volleyball players, and untrained participants (Skurvydas et al., 2011). Marcora et al. showed that PF significantly affected cardiorespiratory responses and exercise performance during high-intensity cycling in healthy men undergoing regular aerobic exercise (Marcora et al., 2008). PF is regulated by a dual system balancing facilitation and inhibition, involving key brain regions such as the limbic system, basal ganglia, motor cortex, dorsolateral prefrontal cortex, and anterior cingulate cortex (Tanaka et al., 2014; Tanaka & Watanabe, 2011). In the context of PF, the neural excitability of the motor cortex decreases, thereby affecting neuromuscular performance (Thomas et al., 2017). This implies that PF may reduce motor cortex activation, thereby leading to a decline in physical performance among high-level athletes.

For the MF condition in the present study, the results showed that MF did not influence jumping, agility, or spiking performance among high-level volleyball players compared with the CG condition. This was in line with a study by Kosack et al., which noted that MF induced by a 60-min Stroop task did not significantly affect CMJ height or badminton performance (Kosack et al., 2020). Holgado et al. found that MF induced by a 90-min AX-continuous performance test did not affect endurance in adult men (Holgado et al., 2021), while Staiano et al. found that MF impaired repeated sprint and jump performance in team sport athletes(Staiano et al., 2023). MF is regulated by an interplay of the limbic system, basal ganglia, frontal cortex, and posterior cingulate cortex (Ishii et al., 2014). MF can affect the activation of the prefrontal cortex, which is an important component of motor networks and plays a role in inhibitory control and fatigue during exercise. However, previous studies have also reported that corticospinal excitability and neuromuscular parameters may not be significantly affected following MF (Holgado et al., 2023; Morris & Christie, 2020; Pageaux et al., 2015). A recent review suggested that nearly half of studies reported no significant effects of MF on various measures of sports performance (Pageaux & Lepers, 2018). This may stem from differences in brain function and motor performance among different individuals. High-level athletes often contend with diverse psychological pressures, including injuries, errors, and negative feedback from spectators, coaches, teammates, and interpersonal conflicts, competition, and rewards (Thompson et al., 2021), while also possessing higher cognitive functions. These findings suggest that isolated MF may not significantly affect motor cortex excitability and neuromuscular capacity among high-level volleyball players.

These results suggested that the combination of physical and mental loads has detrimental effects on jumping, agility, and spiking performance compared with MF alone and the CG condition. Notably, DF led to significantly decreased jumping, agility, and spiking performance at Post 1 and decreased jumping performance at Post 2 compared with PF. This observation aligns with previous research that suggested that DF exacerbates the decline in physical performance more than PF alone (Barzegarpoor et al., 2020; Fuentes-García et al., 2021). Studies have indicated that DF leads to heightened inhibition of prefrontal cortex activity (Mehta & Parasuraman, 2014) and accelerates changes in electroencephalogram activity, resulting in a faster onset of fatigue compared to single-type fatigue conditions (R. Xu et al., 2018). This implies that the inhibition of prefrontal cortex activation due to additional mental activity in the DF condition can potentially affect physical performance in high-level volleyball players.

Our observation indicated that DF and PF did not significantly affect action anticipation in high-level volleyball players when compared with the CG condition. These findings diverge from a study by Alder et al., which reported that individual fatigue types (PF or MF) resulted in decreased accuracy in an anticipation task, with a more pronounced decrease observed for DF, in soccer players (D. B. Alder et al., 2019). In previous studies investigating the effects of fatigue on sport-specific cognitive function, results have not been consistent. Previous studies have reported that PF impaired athletes’ cognitive performance (D. B. Alder et al., 2019; Blain et al., 2019; Parkin et al., 2017) or made top athletes (6 Olympic medalists) perform better in decision-making tasks (Parkin et al., 2017). Inconsistent findings could be attributed to significant variations in the characteristics of the participants (Russell et al., 2019). Action anticipation is an information-processing process that involves visual search, experience, and decision-making (Mann et al., 2010). Long-term sports training can lead to the development of a specialized network for anticipation skills (H. Xu et al., 2016). Thus, fatigue may not significantly disrupt this efficient and focused functional brain connectivity. Fortes et al. reported that MF induced by 30 min of social media use negatively affected decision-making in young male volleyball athletes (Fortes et al., 2021). High-level athletes, owing to their extensive specialized training, have developed a heightened resistance to fatigue (Rubio-Morales et al., 2022) and show better action anticipation during and after exercise compared with at rest (De Waelle et al., 2021; McMorris & Graydon, 1997). This observation implies that high-level volleyball players possess greater fatigue resistance in action anticipation after a prolonged period of specialized training.

We found that DF and MF led to significantly decreased reaction times in the visual search test at Post 2 compared with CG. A recent study compared the effects of 30-min cognitive (Stroop task), physical (65%–75% of max HR cycling), and combined tasks on RT using the psychomotor vigilance task in physically active university students and found that only the MF protocol impaired reaction times (Rubio-Morales et al., 2022). This observation aligns with previous research that suggested that MF led to decreased general cognitive function (Boksem et al., 2005; Duncan et al., 2015; Faber et al., 2012). We also found that only MF significantly decreased reaction times in the anticipation test at Post 1 compared with CG. High-level athletes are often in a state of DF, rather than MF alone in competitive sports. Exercise increases cerebral blood flow and activates brain functional connectivity (Sie et al., 2019; Weinstein et al., 2012), and may counteract the inhibition of brain function caused by MF. Athletes have specialized networks for anticipation in sports scenarios (H. Xu et al., 2016), implying that DF and sports-specific cognitive tasks should be considered when investigating the effects of sports fatigue on athletic performance to improve the ecological validity of future studies.

### Limitations

Despite the use of counterbalancing in the present study, it must be acknowledged that conducting multiple measurements could potentially induce learning effects during the action anticipation task and visual search test. Additionally, fatigue-induced tasks lack high ecological validity. Russel et al. reported that current research lacks perceptual–cognitive sport-specific tasks that can help us better understand the implications of MF on high-level sporting performance (Russell et al., 2019). The conclusions of this study are based on the experimental scenario, and their applicability to real matches or training settings still requires further verification. Sport-based tasks should be used to induce fatigue in future studies. The present investigations focused on the immediate and short-term consequences of fatigue, neglecting potential enduring or long-term effects. It is essential for future longitudinal studies to explore the prolonged effects of fatigue on athletic performance. The absence of notable differences in action anticipation may indicate the presence of unexamined variables influencing performance. Subsequent research should incorporate neurophysiological tools such as electroencephalography and functional magnetic resonance imaging to explore the neurological mechanisms underlying fatigue and its influence on both cognitive and physical performance (Tran et al., 2020). The current study’s investigation into action anticipation aligned with the cognitive skills that neurofeedback training aims to enhance, including concentration, attention, and decision-making—all vital for peak sports performance. The current findings regarding the effects of different forms of fatigue on cognitive function have practical applications in designing neurofeedback protocols. By customizing these protocols, we could enhance athletes’ abilities to withstand MF, thereby potentially improving their predictive skills during competition. This approach underscores the importance of neurofeedback in boosting key cognitive skills for athletes and points to a strategic direction for future research aimed at mitigating the detrimental effects of fatigue on performance (Cheng & Hung, 2020).

### Conclusions

The current study demonstrated that both DF and PF markedly impaired physical performance metrics in elite volleyball players. Notably, the CMJ height, Agility T-test completion time, and spiking performance deteriorated more severely under the DF conditions than under PF, suggesting that the cognitive load in DF may amplify physical performance decrements. This underscores the need for strategies that reduce MF during both training and competitive play. This study thus advocates for the adoption of comprehensive intervention programs aimed at bolstering athletes’ mental endurance, thereby mitigating the adverse effects of MF on physical performance and maintaining cognitive acuity for anticipatory tasks in sports. Anticipation performance was impaired with MF, and visual search behavior was impaired with both MF and DF. This suggested that high-level athletes can counteract the effects of fatigue on anticipation performance. The use of DF and sports-specific cognitive tasks should be considered when investigating the effects of sports fatigue on athletic performance to improve the ecological validity of future studies.

## Authors’ contributions

Y.Y. and F.Q. contributed to the conception and design. Y.Y., M.Z. and Z.L. contributed to data collection and data extraction. Y.Y. and F.Q. drafted the paper. F.Q., M.Z., M.Y.C., and L.Z. revised it critically for important intellectual content. All authors have read and approved the final version of the manuscript, and agree with the order of the presentation of the authors.

